# Gene drive designs for efficient and localisable population suppression using Y-linked editors

**DOI:** 10.1101/2022.06.29.498122

**Authors:** René Geci, Katie Willis, Austin Burt

## Abstract

The sterile insect technique (SIT) has been successful in controlling some pest species but is not practicable for many others due to the large numbers of individuals that need to be reared and released. Previous computer modelling has demonstrated that the release of males carrying a Y-linked editor that kills or sterilises female descendants could be orders of magnitude more efficient than SIT while still remaining spatially restricted, particularly if combined with an autosomal sex distorter. In principle, further gains in efficiency could be achieved by using a self-propagating double drive design, in which each of the two components (the Y-linked editor and the sex ratio distorter) boosted the transmission of the other. To better understand the expected dynamics and impact of releasing constructs of this new design, we have analysed a deterministic population genetic and population dynamic model. Our modelling demonstrates that this design can suppress a population from very low release rates, with no invasion threshold. Importantly, the design can work even if homing rates are low and sex chromosomes are silenced at meiosis, potentially expanding the range of species amenable to such control. Moreover, the predicted dynamics and impacts can be exquisitely sensitive to relatively small (e.g., 25%) changes in allele frequencies in the target population, which could be exploited for sequence-based population targeting. Analysis of published *Anopheles gambiae* genome sequences indicates that even for weakly differentiated populations with an F_ST_ of 0.02 there may be thousands of suitably differentiated genomic sites that could be used to restrict the spread and impact of a release. Our proposed design, which extends an already promising development pathway based on Y-linked editors, is therefore a potentially useful addition to the menu of options for genetic biocontrol.

## Introduction

The sterile insect technique (SIT) of pest control involves the mass rearing, sterilisation, and release of insects that will mate with members of a target population and thereby interfere with their reproduction and reduce their numbers in the next generation (Dyck et al. 2021). The approach has been used with notable success against a number of agricultural pests, including screwworm, fruit flies, pink bollworm and others (Klassen et al. 2021). Sterility has traditionally been achieved by irradiation, but there are closely related approaches using endosymbionts like *Wolbachia* or transgenes conferring dominant lethality that are being explored in mosquitoes and tsetse flies (Bourtzis et al. 2016). It is species-specific in its impacts and becomes more efficient as the target population is suppressed, but rearing and releasing a sufficient number of sufficiently vigorous sterile males to ensure that they mate the majority of females in the target population may not be possible if the insect is difficult to rear or the target population very large, limiting the range of use cases for this otherwise attractive technology.

This requirement for mass releases arises because the sterilising property of the released males disappears when those males die, and every generation a fresh set of sterile males needs to be released. In principle, if a system could be devised in which females are genetically sterilised (or killed), but the genes responsible did not disappear, it could be much more efficient. Burt and Deredec (2018) propose and analyse one way to achieve this objective, which is to use genome editors that are located on the Y chromosome and act in the male germline to make dominant edits on the X chromosome causing those that inherit the X (i.e., daughters) to die or be sterile. The system can also work if the Y-linked editor (YLE) makes dominant edits at an autosomal locus that affect females but not males. In either case the YLE is not itself selected against due to the mutations it creates, because the Y is not found in those females. Thus repeated releases in successive generations can lead to a gradual increase in the frequency of the YLE and the reproductive load imposed on the population, rather than starting from zero each generation, as is the case with sterile males. Computer modelling suggested that this approach could be much more efficient than sterile male releases, requiring perhaps 10-fold fewer males be released. Moreover, the modelling also showed that further efficiencies could be gained by releasing the YLE along with an autosomal sex distorter (ASD) that favours transmission of the Y (e.g., by shredding the X chromosome during male meiosis; Galizi et al. 2014). When released together in the same males, the ASD boosts the transmission rate of the YLE into the next generation, mimicking the effect of a larger release. The ASD itself does not drive, instead being inherited in a Mendelian manner, and would gradually disappear from the population, and so the impacts would still be self-limiting. With this combined system a single release equivalent to less than 10% of the target population may be enough to substantially suppress it, about 100x more efficient than SIT.

These efficiency gains over traditional SIT would be sufficient for many use cases, but in others the release rates required may still be beyond what is feasible in some species, particularly those that are especially difficult to rear and where the target population is particularly large. These cases may require an even more efficient approach. Putting a sex distorter directly onto the Y chromosome, to create a Driving Y, could, in principle, result in a very efficient control system requiring only trivially small releases (Hamilton 1967; Deredec et al. 2008; Deredec et al. 2011). Instead of disrupting the survival of progeny inheriting the X, as a YLE would, such a construct would disrupt the transmission of the X chromosome, thereby favouring its own transmission. All else being equal, the Driving Y would increase in frequency over successive generations, gradually making the sex ratio more and more male biased, and thereby suppressing or even eliminating the population. However, while there are many natural examples of driving sex chromosomes (Jaenike 2001; Burt and Trivers 2006), including driving male-determining regions in two species of mosquito (Wood and Newton 1991), and sex chromosome distorters have been engineered that work if inserted on an autosome (Galizi et al. 2014; Galizi et al. 2016; Fasulo et al. 2020; Meccariello et al. 2021), synthetic Driving Ys are thus far proving difficult to engineer, due at least in part to the silencing of the sex chromosomes at male meiosis seen in many species (Taxiarchi et al. 2019).

In this paper we return to the idea of having separate YLE and ASD constructs, and explore the consequences of making a small addition to the autosomal construct that allows not only the ASD to increase the transmission of the YLE, but also for the YLE to increase the transmission of the ASD, to make a “double drive” (Willis and Burt 2021). In particular, we consider a design in which a gRNA gene is added to the ASD that allows it to home in the presence of the YLE. As will be seen, this small alteration in the design changes the dynamics considerably, so that trivially small release rates can lead to spread of the constructs and suppression of the population, similar to a Driving Y. Importantly, the approach does not rely on sex chromosome expression during meiosis, and can work even with low homing rates, thereby potentially expanding the range of species for which low-release-rate population suppression gene drive approaches are feasible. Moreover, we show that by appropriate design, relatively small pre-existing differences in allele frequencies at a polymorphic site between target and non-target populations of the same species (or species complex) can be used to restrict the spread and impact of the drives (though if control is desired throughout a species range, that would be possible too). We then analyse published genome sequence data from *Anopheles gambiae* s.l. mosquitoes which suggests even weakly differentiated populations could, in principle, be differentially targeted with appropriately designed constructs. These results demonstrate the potential flexibility of YLEs for population suppression, with an incremental step-by-step development pathway of increasingly efficacious constructs, from self-limiting genome editors to self-sustaining double drives, each step building molecularly on the one before.

## Methods

### Genetic design

The design under consideration is an extension of those proposed and analysed by Burt and Deredec (2018). It consists of two constructs, one on the Y chromosome and one on an autosome. A number of different molecular constructions could be used, but for concreteness we will assume a specific configuration (Fig. 1). The Y-linked construct encodes a genome editor that targets a gene on the X-chromosome, and the edits induced cause dominant lethality or sterility in the progeny. Because the progeny of a male that inherit his X chromosome are female, it is his daughters that are killed or made sterile. In the specific example configuration, the editor consists of a Cas9 nuclease gene with control sequences that mean it gets expressed in the male germline, and a ubiquitously expressed gRNA gene targeting the X-linked target gene. The autosomal construct encodes a nuclease that targets a repeated sequence on the X chromosome (separate from the target of the YLE), thereby shredding the X, and is expressed during male meiosis, so that males with the construct produce mostly Y-bearing sperm and male offspring (Galizi et al. 2014). It also carries a ubiquitously expressed gRNA gene targeting its insertion site, so that in heterozygous males, in the presence of the Y-linked construct, the autosomal construct can home in the germline and thereby be transmitted at a greater than Mendelian rate to the offspring. It is the addition of this autosomal gRNA gene that distinguishes this design from the “augmented YLE” design modelled by Burt and Deredec (2018) and the autosomal sex distorter strains published by Galizi et al (2014).

**Figure 1:**
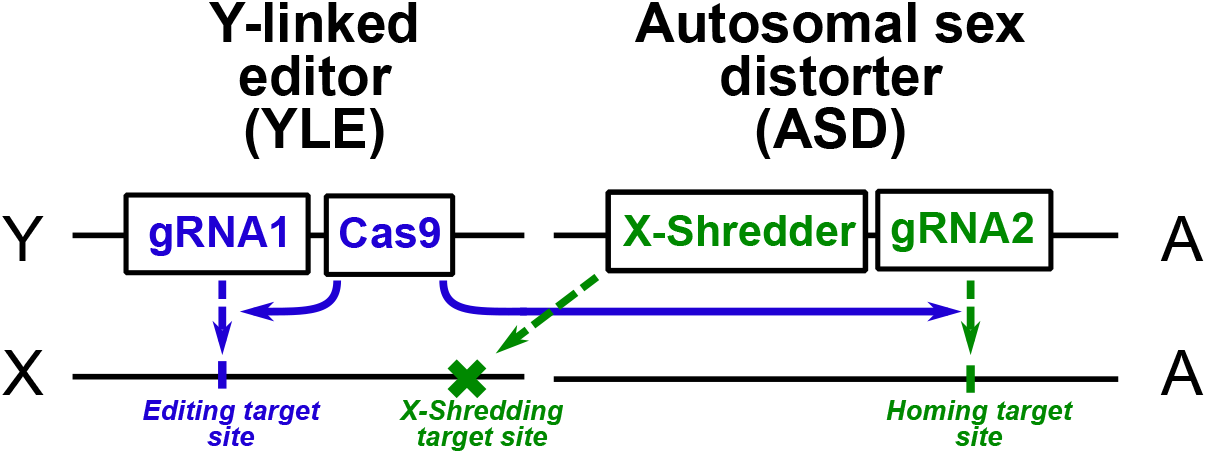
The proposed genetic constructs. We model a Y-linked editor (YLE) that targets an X-linked locus such that female offspring die and an autosomal sex distorter (ASD) that in males disrupts transmission of the X chromosome (thereby favouring transmission of the Y), and that, in the presence of the YLE, shows drive via the homing reaction. More specifically, we model the YLE as consisting of a germline-expressed Cas9 and ubiquitously expressed gRNA (together responsible for the editing), and the ASD as consisting of a X-shredder nuclease expressed during male meiosis (responsible for the sex distortion) and a ubiquitously expressed gRNA that, in the presence of the Y-linked Cas9, allows the ASD to home.

Note that we now have a double drive (Willis and Burt 2021) in which the transmission of each construct depends in some way on the presence of the other, though in somewhat different ways. The X-shredder causes whatever Y chromosome it is with to show superMendelian transmission, and if there is a statistical association (i.e., linkage disequilibrium) between the two constructs, then the YLE will increase in frequency due to the presence of the X-shredder. At the same time, the autosomal construct can show superMendelian transmission via the homing reaction, but only in the presence of the Y-linked construct. Note also that the editor function *per se* is not directly involved in the drive of either construct, and even if the gRNA responsible for the editing was absent the rest would act as a double drive and lead to some suppression of the target population due to the sex ratio distortion produced by the X-shredder. However, as we will see, including the editor function makes a key quantitative contribution to the function of the whole in terms of increasing the magnitude and duration of suppression.

### Modelling

We use a deterministic population genetic and population dynamic model based on one used previously (Burt and Deredec 2018; Willis and Burt 2021) to investigate the fate of the proposed genetic constructs and their impacts on population size. In brief, the model assumes non-overlapping generations and two life stages, juveniles and adults, with density-dependent mortality occurring during the juvenile phase. Mating is random, and all females are assumed to mate (i.e., males are not limiting). The constructs are introduced into a target population by release of heterozygous males equivalent to 0.1% of the target population.

The model allows for both intended and unintended fitness effects. The intended fitness effects include the effect of the edited allele on female fitness (our baseline assumption is it causes fully dominant and penetrant lethality of adult females, after density-dependent mortality and before censusing) and the fitness effects that arise automatically from distortions of the sex ratio (Shaw and Mohler 1953). Unintended fitness costs include those due to expression of the proteins and gRNAs (e.g., the energetic costs of their synthesis) and potential costs due to the activity of the various editors and nucleases (e.g., due to off-target events elsewhere in the genome).

Finally, our model considers a variety of types of mutation. Each component of each construct can be affected by loss-of-function mutations occurring at a constant background mutation rate. Loss-of-function mutations can also occur on the autosomal construct at an elevated rate during homing.

Finally, the processes of editing, shredding, and homing can lead to the production of functional resistant sequences at each of the three target sites (e.g., through end joining repair). For simplicity we assume the resistant sequences have the same fitness as the wild-type alleles. More details about the mechanics of the model and a table of parameters and their baseline values are given in the Supplementary Information. Model code and simulation results are freely available on GitHub (https://github.com/ReneGeci/LocalisableYLEsuppression).

### *An. gambiae* PAM site analysis

To better understand the feasibility of differentially targeting populations of the same species we analysed published genomic sequence from 15 populations of the malaria mosquito *An. gambiae* (The *Anopheles gambiae* 1000 Genomes Consortium 2020). The most commonly used CRISPR-Cas9 system requires an NGG sequence in the protospacer adjacent motif (PAM), so we screened for polymorphic GG (or CC) dinucleotide sites and for each pair of populations scored the number of sites that showed the appropriate difference in allele frequencies as determined by the modelling. Further details are given in the Supplementary Information.

## Results

### Idealised case

We first consider the simplest idealised case where there are no unintended fitness costs, loss-of-function mutations, or functional or non-functional resistance, and assume molecular efficiencies (editing, homing, X-shredding rates) are high but not perfect, within the range seen in *An. gambiae* mosquitoes (90-95%; Galizi et al. 2014; Hammond et al. 2016; Kyrou et al. 2018; Carballar-Lejarazu et al. 2020; Simoni et al. 2020). In this case if males carrying the two constructs are released, then both the YLE and the ASD constructs increase to and remain at a very high frequency (Fig. 2a). The YLE increases in frequency because of the statistical association between it and the ASD. Initially, this correlation is 1 (because they are released in the same males), but gradually it declines because of less than perfect homing and X-shredding rates. The ASD increases in frequency because it homes in the presence of the YLE. It does not home in females or in wildtype males and spreads more slowly than the YLE. As a result of the joint action of the YLE causing dominant female sterile mutations and the ASD producing a male-biased sex ratio, the number of females (and therefore total population size) crashes in about 20 generations.

**Fig.2.**
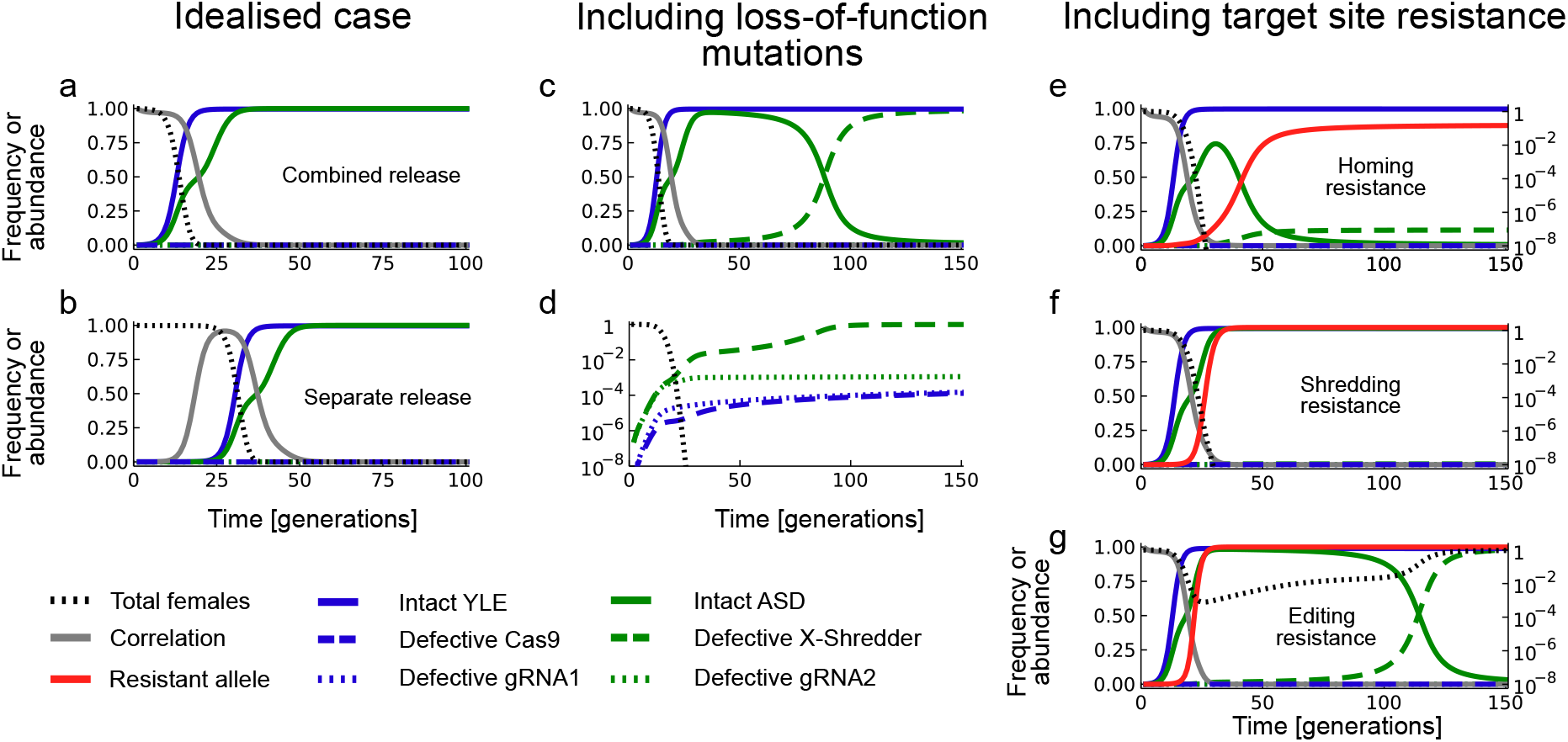
Timecourse of gene and population dynamics. (a, b) Idealised case (no mutation, no resistance, no unintended fitness costs) with release of constructs in the same males (a) or in separate males (b). (c, d) Allowing for loss-of-function mutations in the constructs, with the frequency of the most common alleles on an arithmetic scale (c) and frequency of defective variants on a log scale (d). (e-g) Allowing for end-joining repair to produce resistant alleles at the site of homing (e), shredding (f) or editing (g). Allele frequencies are shown on the arithmetic scale (left) and population size (number of females) on the log scale (right). All population sizes are relative to the pre-release equilibrium.

Though the spread of the YLE depends on a statistical correlation with the ASD, it is not necessary for the two constructs to be released in the same males for them to spread. If they are released in different males (in which case the correlation between them is initially negative), the fact that the YLE helps the ASD to drive means that a positive correlation gradually builds up, allowing the two constructs to spread, before the correlation falls again (Fig. 2b). The dynamics of spread and impact on the population are much the same as with combined releases, just with a delay. On the other hand, neither construct, if released by itself, increases in frequency (Supplementary Figure SF-1).

### Including loss-of-function mutations

In the idealised model just considered there are two types of Y (wildtype or transgenic), two types of X (wildtype or edited), and two types of autosome (wildtype or transgenic). We now consider the more realistic case in which mutations can arise in the various components of the two constructs which abolishes their function. We assume the loss-of-function mutations occur in the two components of the YLE (the Cas9 and the gRNA) and the two components of the ASD (the X-shredder and the gRNA) with a probability of 1e-6 per generation, due to errors of normal DNA replication, and, moreover, that mutations arise in the two components of the autosomal construct with a probability of 1e-3 per homing event, as homing is likely to be associated with a larger mutation rate than normal DNA replication (Hicks et al. 2010; Simoni et al. 2014; Rodgers and McVey 2016).

The dynamics in this case are initially the same as in the idealised case, with the two constructs increasing in frequency, and population size declining (Fig. 2c). But then a defective autosomal construct that has the gRNA but is missing the X-shredder begins to accumulate, replacing the intact construct. These defective elements arise predominantly during the homing reaction, and they rapidly increase in frequency because they are still able to home and yet are associated with a 50:50 sex ratio, which is strongly selected for in a population that has become strongly male-biased. Moreover, the correlation between the YLE and ASD is small by this time, so X-shredding has less of an effect on the probability of homing in the next generation than immediately after the release.

Despite the loss of the X-shredder, the population is still very substantially suppressed, because the YLE is still very frequent: the X-shredder functions primarily as a booster of the YLE, not as a source of reproductive load itself.

Defective variants of the other three components of the original constructs do not have the same dynamics and remain rare (Fig. 2d). The gRNA in the autosomal construct allows it to home, and so loss-of-function mutations in it are selected against as long as there are still sites to home in to. The Cas9 in the YLE helps maintain the association with the X-shredder, and so loss of that too will be selected against. Finally, the gRNA in the YLE does not help the YLE spread, but the pressure on it to be lost is weak. Since this construct does not home, the only source of loss-of-function mutations is normal DNA replication, and these arise rarely (1e-6 in our model) and accumulate slowly. If there are fitness costs associated with any of these functions (e.g., due to costs of gRNA or Cas9 synthesis, or off-target cleavage), then these might accelerate the loss of one or other of these functions, but the effect will be modest unless the costs are large.

### Target site resistance

The three molecular processes in our design (editing, shredding, and homing) each occur at a different target site, and each one could give rise to a resistant sequence that is no longer recognised by the enzymes involved (e.g., by end-joining repair, EJR). We now consider the consequences of allowing such mutations to arise, assuming the most conservative case where resistant alleles have the same fitness as the wildtype allele.

#### Homing resistance

The insertion site of the ASD is not specified in Fig. 1 and our baseline assumption is that the insertion has no fitness cost. The simplest way to achieve this would be to insert it in a neutral non-functional part of genome, in which case all cleavage-resistant sequences are as “functional” as the wildtype, and the frequency of functional resistance mutations is then just the frequency of end-joining repair (plus any partial homing events producing wholly defective inserts). The effect of including even a small frequency of EJR (5%) at the autosomal locus is that it is no longer the defective X-shredder that accumulates, but instead the resistant allele (Fig, 2e).

Nevertheless, we still observe strong suppression, even with EJR rates of up to 75% (Fig. 3a). Again, the spread of the ASD is not directly responsible for suppressing the population – it merely functions to boost the YLE to a high frequency, and it does not have to reach a high frequency itself to achieve that. This ability of the design to function even in species with weak and inefficient homing rates could be an attractive feature, expanding the range of species in which such control is possible, and, because of the potential interest, all subsequent modelling in this paper will be done for both low (5%) and high (60%) EJR scenarios or species.

**Fig. 3.**
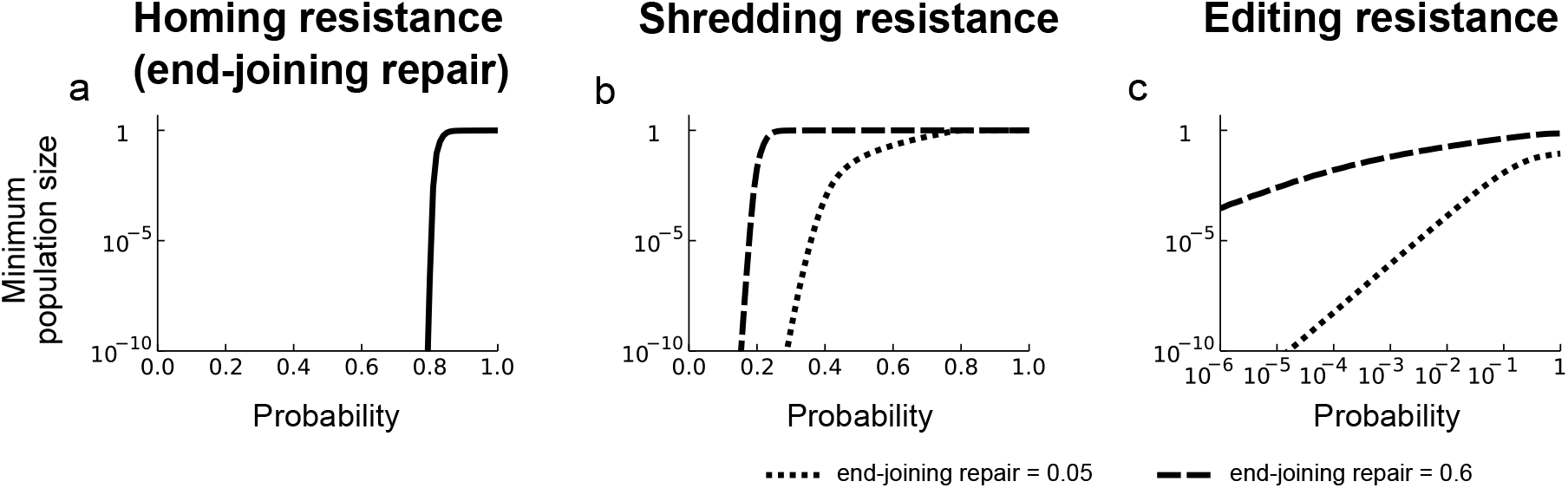
The impact of target site resistance. Minimum population size (relative to the pre-release equilibrium) as a function of the probability of a resistant allele forming at the site of homing (a), shredding (b), and editing (c; note log X-axis). For (b) and (c) results are shown for both low and high EJR scenarios (5% and 60% EJR, respectively).

#### Shredding resistance

The X-shredder is likely to need to target a repeated sequence on the X, and so it may be that resistance would require simultaneous changes at many sequences and may therefore be unlikely to arise. Nevertheless, we can still assess the consequences if it did. If we assume it arises in 5% of shredding events, then the modelling again shows that resistance evolves quickly but the population is still very substantially suppressed (Fig. 2f). Again, it is not the shredding that is primarily responsible for the load on the population, but the editing. Even higher probabilities of shredding resistance arising can still lead to substantial suppression, even in high EJR species (Fig. 3b). Note that if shredding resistance evolves then there is no selection in favour of the defective X-shredder element, because shredding is not occurring anyway (Fig. 2f).

#### Editing resistance

Finally, if we allow for editing resistance at 5%, then, again, resistance evolves very quickly, but now the population is suppressed to a far lesser extent (minimum population size 8e-4 relative to the original, after which it recovers; Fig. 2g). Moreover, while for both homing and shredding resistance there is a relatively rapid transition between very strong suppression and virtually none as the probabilities of resistance arising change, for editing resistance the picture is somewhat different (Fig. 3c). As noted above, even if there is no editing at all, the double drive can lead to some level of suppression due to the sex-distorting effect of the X-shredder. However, most of the load imposed on a population is due to the editor, and if functional resistance does arise at the target of editing it is rapidly selected for. As a result, even low rates of resistance (1e-3 or 1e-2)^1^ significantly reduces the level of suppression. This is particularly the case in high EJR species; the strategy is somewhat more resilient in low EJR species. Nevertheless, the modelling suggests that most effort and attention in preventing functional resistance should go into the design of the editing function.

### Sensitivity analysis for other parameters

The model includes a number of other parameters, and the impact of varying these (while leaving the others at their baseline values) is shown in Fig. 4, for both low and high EJR scenarios. The effect of varying the efficiencies of the three processes is as expected from the resistance analysis: the requirements on homing efficiencies are the least stringent, followed by the shredding efficiency, and finally the requirement on editing efficiency is the most stringent, with rates needing to be greater than about 80% in high EJR species to have a strong suppressive effect (Fig. 4a-c). Note again that if editing efficiency is 0 there can still be some suppression, due to the sex ratio distortion produced by the X-shredder, but it is relatively small (Fig. 4c). The effect of the edit on female fitness must be similarly high and dominant (Fig. 4d-e).

**Fig. 4.**
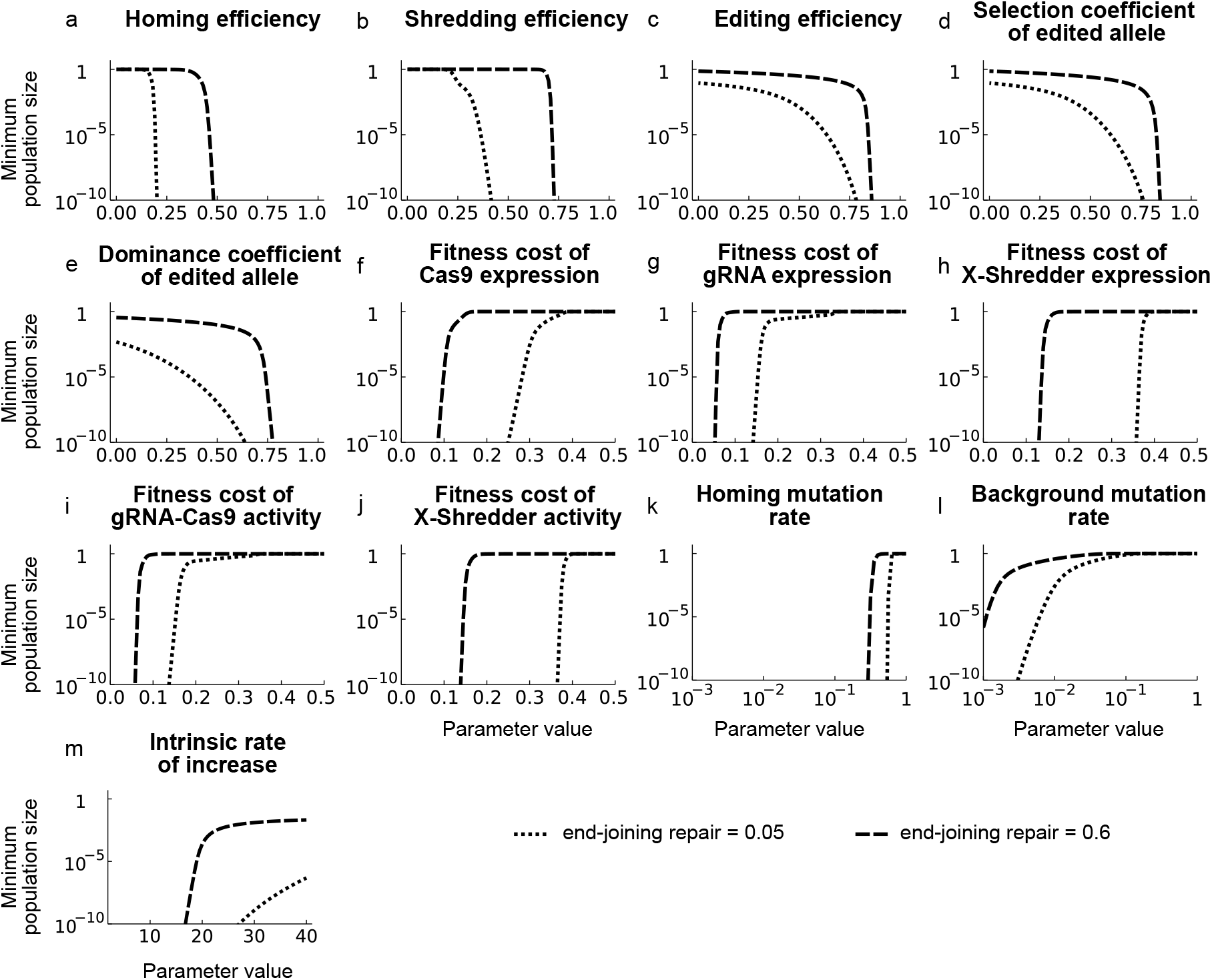
Sensitivity analysis. Each plot shows the minimum population size as a function of changes in the specified parameter assuming low EJR (5%; dotted lines) or high (60%; solid lines). Note for (a) that the homing rate is equal to (cleavage rate) x (1-end joining rate).

Six unintended fitness costs are modelled. Under our baseline assumptions the fitness cost of the edited allele in males does not matter because the edited allele never occurs in males: it is created on the X-chromosome in the male germline, and then transmitted to daughters, who die with 100% probability. If the target of editing was autosomal, then fitness effects in males would matter more (Burt and Deredec 2018). For the other five unintended fitness costs there is a relatively sharp threshold between very good suppression and virtually none somewhere between fitness costs of about 0.15 and 0.35 (low EJR), or 0.05 and 0.15 (high EJR) (Fig. 4f-j). The requirements on costs of gRNA expression and of Cas9-gRNA activity are the most stringent, presumably because each of them is doubled due to there being two gRNAs.

Population suppression is robust to reasonable values for loss-of-function mutation rates. For homing-associated mutations, rates up to about 50% are consistent with strong suppression, and for normal cell-division-associated mutations rates of up to 1e-3 (Fig. 4k-l). The rate of recombination in females between the two X-linked loci, the target of X-shredding and the target of editing, has no significant effect because, under the baseline assumptions, females with a single copy of the edited allele are dominant lethal. Finally, population suppression is also robust to reasonable values for the intrinsic rate of population increase (Fig. 4m).

### Population restriction using allele frequency differences

The ability of the double drive to increase in frequency from rare means that it will spread not only in the release population, but also in other populations into which there is even a small level of gene flow. This ability to spread across a landscape can contribute greatly to the efficiency of the release, but may also cause problems, as there may be non-target populations one would not want to impact. In principle, one way to restrict the spread of a gene drive is to exploit pre-existing sequence differences between target and non-target populations (Sudweeks et al. 2019; Willis and Burt 2021).

We now investigate how such targeting would work for our proposed design by considering the impact of different levels of pre-existing resistance.

As noted above, 100% editing resistance can still lead to the spread of the constructs and some suppression, so choosing a site for editing that is differentiated between target and non-target populations will not provide effective restriction. For the other two processes, homing and shredding, we simulated releases into populations with different levels of pre-existing resistance (example time courses shown in Supplementary Figure SF-2). Plots showing the maximum construct frequencies and minimum population sizes as a function of the frequency of pre-existing resistance for low and high EJR scenarios are shown in Figure 5. In each of the four cases analysed there is good suppression with no pre-existing resistance, and negligible suppression with 100% pre-existing resistance. For homing resistance in low EJR species the transition between these occurs sharply, with pre-existing resistance of 84% giving a minimum population size of 1e-6 (i.e., 99.9999% suppression), and pre-existing resistance of 87% giving a minimum population size of 0.95 (i.e., 5% suppression). However, suppression in the former case can take impracticably long (hundreds of generations); if we specify that the target population must be suppressed by 99% within 50 generations, then that requires the pre-existing resistant allele have a frequency less than 64%.

**Fig. 5.**
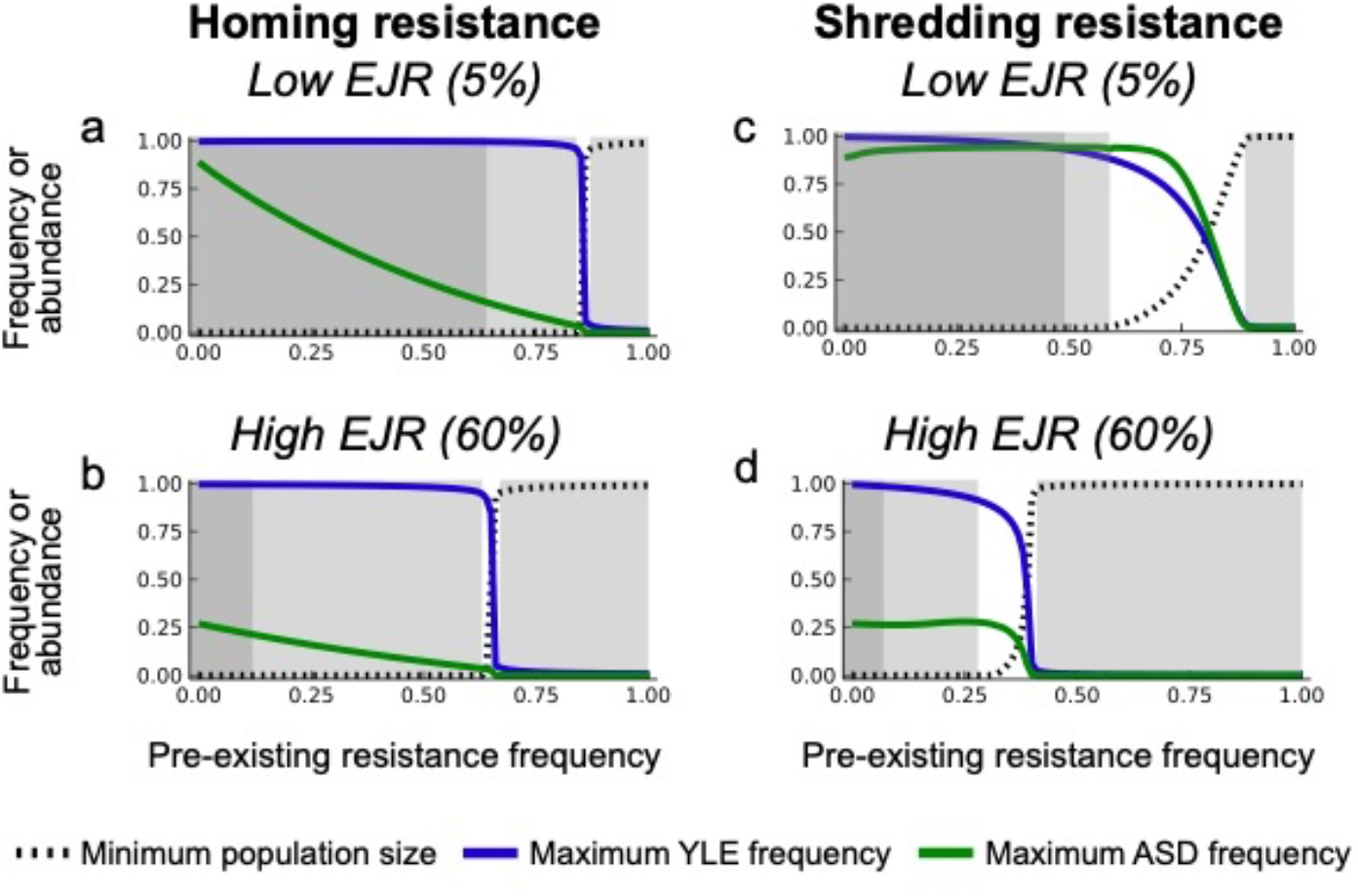
Exploiting pre-existing sequence differences to restrict population impact. Minimum population size and maximum construct frequencies as a function of the frequency of pre-existing resistance at the sites of homing and shredding for low and high EJR scenarios. Simulations were run for 1000 generations with different initial frequencies of resistant alleles. In each plot the lightly shaded boxes to the left and right show regions where the minimum population size over the 1000 generations is less than 1e-6 or greater than 0.95, respectively, as would be appropriate for target and non-target populations, and the darkly shaded box to the left shows regions where the population is suppressed by at least 99% in 50 generations, and therefore control may be practicable.

From an engineering point of view the fact that this transition from substantial to negligible control occurs in a relatively small window of pre-existing sequence differences (64—87%) is a good thing, as it reduces the level of overall population differentiation needed to find a suitably differentiated target site that would restrict the impact of a release to target populations. In the ideal case it would resemble a step function. The transition is less sharp for homing resistance in high EJR species (12— 67%), and for shredding resistance in low and high EJR species the transition is intermediate between these extremes (Fig. 5).

### PAM site analysis in *An. gambiae*

To better understand the potential opportunities for sequence-based population targeting that these results imply, we performed a preliminary analysis of published genome sequences from *An. gambiae* s.l. mosquitoes, analysing data from 15 populations and 1138 individuals across sub-Saharan Africa (The *Anopheles gambiae* 1000 Genomes Consortium 2020). The most commonly used Cas9-based CRISPR nuclease recognises a protospacer adjacent motif (PAM) of NGG, so we screened the database for polymorphic GG (or CC) dinucleotides. After filtering out sites with more than 5% missing data within any population, we were left with 13,462,450 polymorphic PAMs. For each pair of populations we counted the number of PAM sites that had a frequency in one population >36% (the target population) and in the other <13% (non-target population), representing the level of differentiation required at the homing site in a low EJR species (fig 5a), and then re-did the count in the opposite direction (i.e., reversing the assignments of target and non-target population). The average for the two directions was then plotted against the F_ST_ for that pair of populations, the standard overall measure of population differentiation (Fig. 6). As expected, the number of suitable PAM sites increased with F_ST_, and for all population pairs with F_ST_ > 0.02 there were more than 4000 appropriately differentiated PAM sites. For less well differentiated populations there was substantial variation in the number of suitable PAM sites, from 200 – 8000 sites. We repeated the analysis using the thresholds for high EJR species (>88% and <33%), and, as expected, the number of suitable PAMs was lower, fewer than 10 in some instances, but most population pairs had hundreds to tens of thousands. For extrapolating to other species, we note that the combined length of the autosomes in *An. gambiae* is about 206Mb, of which about 61% was accessible in these analyses.

**Fig. 6.**
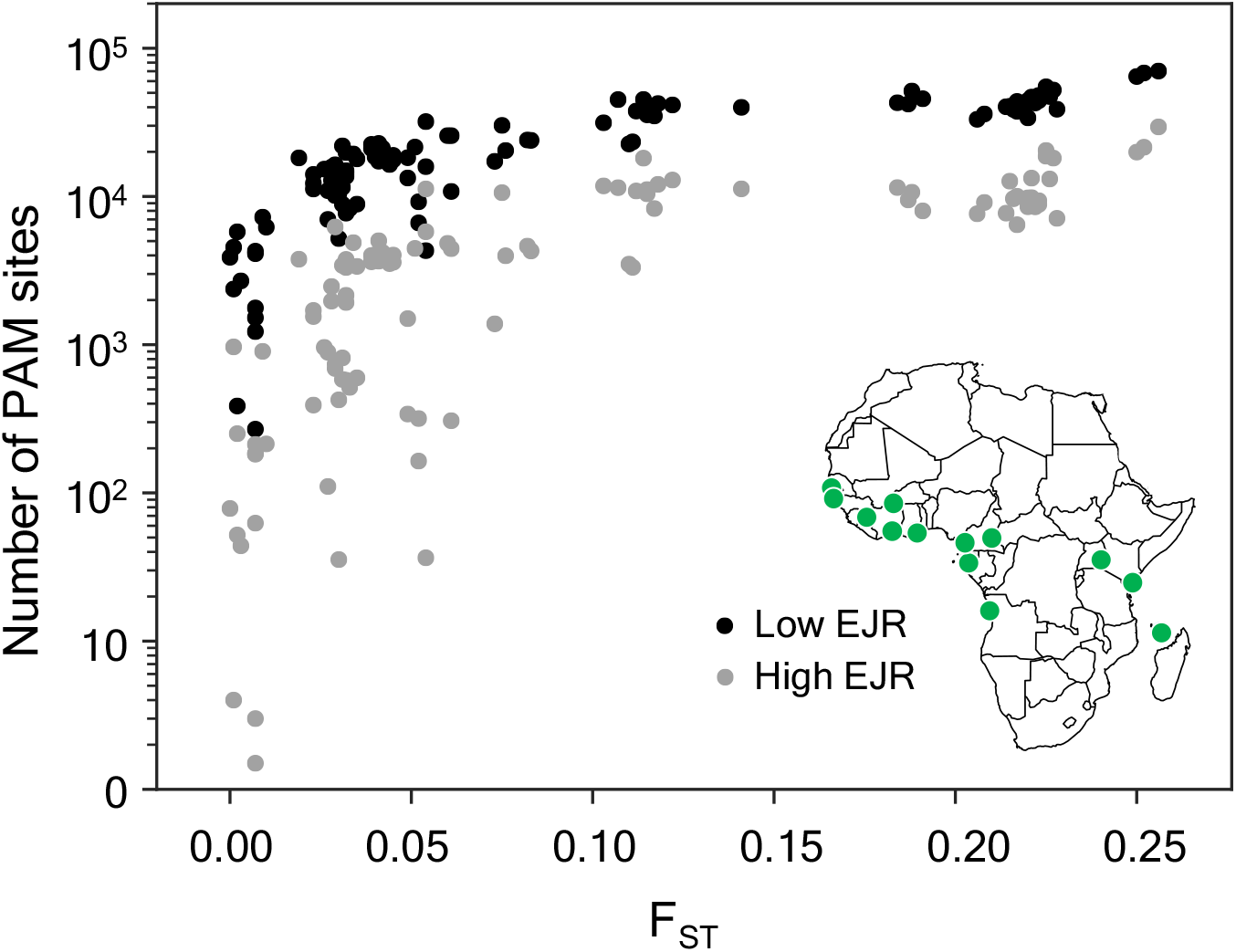
Number of suitably differentiated PAM sites as a function of the overall differentiation between pairs of *An. gambiae* sl populations. For each pair of populations we plot the average number of autosomal PAM sites with frequencies of >36% in one population and <13% in the other (low EJR scenario, black dots, equivalent to frequencies of resistant alleles of <64% and >87%, respectively) or >88% in one population and <33% in the other (high EJR scenario, blue dots). F_ST_ values were obtained from The *Anopheles gambiae* 1000 Genomes Consortium (2020, Supp. Fig. S5). Inset shows the location of the 15 populations.

## Discussion

Burt and Deredec (2018) have demonstrated that YLEs could, in principle, provide highly efficient yet self-limiting population control, either on their own or augmented with an ASD. In principle, substantial suppression could be achieved even with a single release of less than 10% of the target population. However, for some potential use cases, even this number may be prohibitive, and more efficient strategies could be achieved with self-sustaining gene drive designs. In this paper we have proposed and analysed a method for getting the YLE to spread in a self-sustaining way using a double drive design, defined as “one that uses two constructs, inserted at different locations in the genome, both of which can increase in frequency, at least initially, and which interact such that the transmission of at least one of them depends on the other” (Willis and Burt 2021). The specific molecular adjustment made was to add a gRNA to the ASD, which means that not only does the ASD boost the transmission of the YLE, but the YLE also boosts the transmission of the ASD. This reciprocity allows the constructs to spread from rare, even if they are initially released in different males, and so they behave cooperatively to spread selfishly (from the rest of the genome’s point of view). While there is reciprocity, the two constructs are not symmetrical and there is a division of labour between the YLE that is primarily responsible for imposing the reproductive load and the ASD that is responsible for the spread of the first.

Though we have analysed only a well-mixed non-spatial model, it is possible to make predictions about the dynamics and impact of a release across a landscape. The ability of the double drive to spread from rare, even if they are not initially in the same organisms, means that the constructs would be expected to spread geographically, to any population with which there is some reasonable level of gene flow. This is so despite a tendency for the intact ASD to eventually be lost within a population (Fig. 2c, e). As long as the YLE and ASD are rare, the intact ASD is positively selected for, and as long as a few functional types reach a population, spread will ensue, so populations could be suppressed even far from the release site. That is, the X-shredder function should be evolutionarily stable enough to allow spread across an entire landscape. The editing function, on the other hand, though primarily responsible for the reproductive load imposed on the population, does not contribute to the spread of the constructs, and one would expect that eventually, if the species is sufficiently widespread (relative to dispersal distances), the gRNA component of the YLE will be lost. However, if the fitness cost of the gRNA in males is low, and the mutation rate (which only involves normal cell-division-associated DNA replication) is low, there could still be significant geographical suppression before that happened. This dynamic is analogous to a population replacement gene drive losing a cargo effector gene (Beaghton et al. 2017), but here the mutation rate may be much smaller (because the YLE does not home), so, all else being equal, it should persist longer. In principle, it may be possible to engineer YLEs that will remain intact for different distances from the point of release by altering the mutation rate (e.g., adding repeated sequences to increase; adding redundant gRNAs to decrease) or fitness effects, though a reduced double drive will continue to spread beyond the point of the gRNA being lost, having a transient impact on population size due to the X-shredder. As in other population suppression gene drive systems, after the initial spread across a landscape there may be complex extinction-recolonisation or chasing dynamics (Eckhoff et al. 2017; North et al. 2019; North et al. 2020; Champer et al. 2021), particularly in the absence of inbreeding depression or other strong Allee effects (Beaghton and Burt 2022), but further investigation is beyond the scope of this study.

As with any proposed form of pest control, one needs to consider the possibility of resistance evolving. Because the reproductive load imposed on the population is primarily due to the edited locus, selection for target site resistance is strongest at that site, and the extent of population suppression is most sensitive to the probability of resistance arising at this site (Fig. 3). Efforts to reduce the likelihood of resistance evolving should therefore be focussed on this site. This can be done by targeting key functional sites that are not able to change while maintaining function, and by targeting multiple neighbouring sites in the same target gene (Champer et al. 2018; Kyrou et al. 2018; Champer et al. 2020). By comparison, selection for resistance at the sites of homing and shredding is weaker, and higher rates of mutation to resistance can be tolerated.

Similar considerations apply to the required efficiencies of the different processes. Our sensitivity analysis shows that the requirements for the editing efficiency are the most stringent (>80% in the high EJR scenario with our baseline parameter values), while the requirements for the shredding and homing rates are lower. This difference arises because the YLE is insulated from the negative fitness effects it causes to daughters, and therefore selection against it is weak, and the processes increasing its frequency (shredding and homing) do not have to be very strong in order to get the YLE to a high frequency. Indeed, we have seen that the design can work even if most cleavage events at the autosomal site are repaired by end-joining. This robustness may be important because while high rates of homing and shredding have been achieved in anopheline mosquitoes, the rates observed thus far in some other species have been lower (Grunwald et al. 2019; Fasulo et al. 2020; Meccariello et al. 2021). Thus our design may expand the range of species in which it is possible to engineer low release rate population suppression gene drives. We speculate that it may be possible to further reduce the required rates of homing and shredding if the X-shredding activity is conditional, only occurring in the presence of both constructs (Burt and Deredec 2018), though this may be more difficult to engineer if the sex chromosomes are silenced at meiosis. The design analysed here does not rely on expression of the sex chromosomes during meiosis, though it does require expression from the Y-chromosome in the germline. Such expression has been observed for a marker gene in *An. gambiae* (Bernardini et al. 2014) and for Cas9 in *Drosophila melanogaster* (Gamez et al. 2021).

A particularly attractive feature of our design is the potential it offers for localisation of the spread and impact of the releases based on pre-existing sequence differences between target and non-target populations of the same species or species complex. This ability was also found in the autosomal double drives of Willis and Burt (2021), but with the design analysed here even smaller differences in allele frequency can be exploited to localise the spread and impact, as seen in the relatively steep change in outcome based on small differences in allele frequencies (Fig. 5). This sensitive dependence of the dynamics on initial conditions (the frequency of pre-existing resistance) is reminiscent of that seen with so-called threshold-dependent gene drive designs, which spread if introduced above a particular frequency, and disappear if introduced below that frequency, and have been proposed as an alternative method to localise spread and impacts (Dhole et al. 2018; Dhole et al. 2019; de Haas and Otto 2020; Edgington et al. 2020; Greenbaum et al. 2021). Our PAM site analysis of *An. gambiae* population genomic data indicates there may be very many suitably diverged target regions of the genome between even closely related populations. Such sequence-dependent localisation could be used to limit the spread and impact of a release either geographically or taxonomically, whenever the target taxon interbreeds to some extent with non-target taxa. That said, our analysis must be considered preliminary. For example, we have assumed that any difference in PAM site would be enough to block cleavage, but this assumption would need to be tested in the target species. In some systems the Cas9 from *Streptococcus pyogenes* can recognise not only the canonical NGG PAM, but also NAG, NGA, and NNGG, albeit with lower efficiency (Collias and Beisel 2021), and one might need to avoid these polymorphisms if the same occurs in the species of interest.

The final key feature of our design worth emphasising is that it is based on a small molecular modification of highly efficient self-limiting designs. Modern biotechnology has the potential to offer effective and sustainable forms of pest control that are safe for both people and the environment, but the very fact they are new has led to recommendations for a step-by-step approach in which increasingly efficacious constructs are assessed (NASEM 2016; WHO 2021). The starting point for our design is a genome editor that acts in males to produce dominant X-linked lethal or sterile mutations. Some of the molecular options for the editor are outlined in Burt and Deredec (2018), including knocking out a haplo-insufficient gene, producing a dominant negative mutation, or using paternal deposition of the editor, and a proof-of-principle demonstration in *D. melanogaster* is provided by Fasulo et al. (2020). Instead of targeting an X-linked locus, it would also be possible to target one on an autosome, but the impact would have to be female-specific; potential targets are rare but do exist (Navarro-Paya et al. 2020; Willis and Burt 2021). From this starting point, it is now possible to identify a step-wise progression of at least five designs of increasing efficacy:

1. An autosomal insertion of the construct, which would be transmitted 50% of the time to the daughters that are killed, and so halve in frequency each generation, and therefore have dynamics like a female-specific RIDL strain (Gentile et al. 2015; Vella et al. 2021)
2. The same construct moved to the Y-chromosome (i.e., a YLE)
3. Augmenting the releases with an autosomal sex distorter to boost the frequency of the YLE
4. A localised sequence-specific YLE-based double drive (i.e., the design analysed in this paper)
5. A ‘universal’ (non-population-specific) YLE-based double drive

We are not suggesting that it will be necessary or even desirable to go through all five steps for all possible use cases. Rather, we merely wish to note that there is a great deal of flexibility in the designs, and therefore in the development pipeline, which can be optimised for any specific use case. This sort of step-by-step development pathway may prove attractive to developers, regulators, and other stakeholders more generally (Nash et al. 2019).

Sex chromosome systems vary widely among species, and in some species the repetitive and heterochromatic structure of the Y chromosome can make them difficult to work with. However, as we have seen, there can be considerable advantages to developing YLE-based interventions, including highly efficient self-limiting strategies; self-sustaining strategies that can be effective with lower rates of homologous repair and no requirement for meiotic expression off the sex chromosomes; more options for using pre-existing sequence differences to limit the spread and impact of a self-sustaining release; and a step-by-step development pipeline which may help reduce uncertainties and facilitate acceptance by developers, regulators, other stakeholders, and the relevant publics at large. Therefore, despite the difficulties of working with the Y chromosome, given the potential benefits of the approach, it seems well worth investing the time and effort needed to fully explore this approach. The recent rapid expansion of molecular engineering capabilities in multiple species (Hay et al. 2021; Bier 2022) makes us optimistic the challenges can be met.

## Acknowledgements

We thank John Connolly, Silke Fuchs and John Mumford for useful comments on a previous draft. Supported by grants from the Bill & Melinda Gates Foundation (Grant INV006610 “Target Malaria Phase II”) and the Open Philanthropy Project Fund, an advised fund of Silicon Valley Community Foundation (Grant O-77157) to AB. The funders had no role in study design, data collection and analysis, decision to publish, or preparation of the manuscript.

## Supplementary figures

**Supplementary Figure SF-1:**
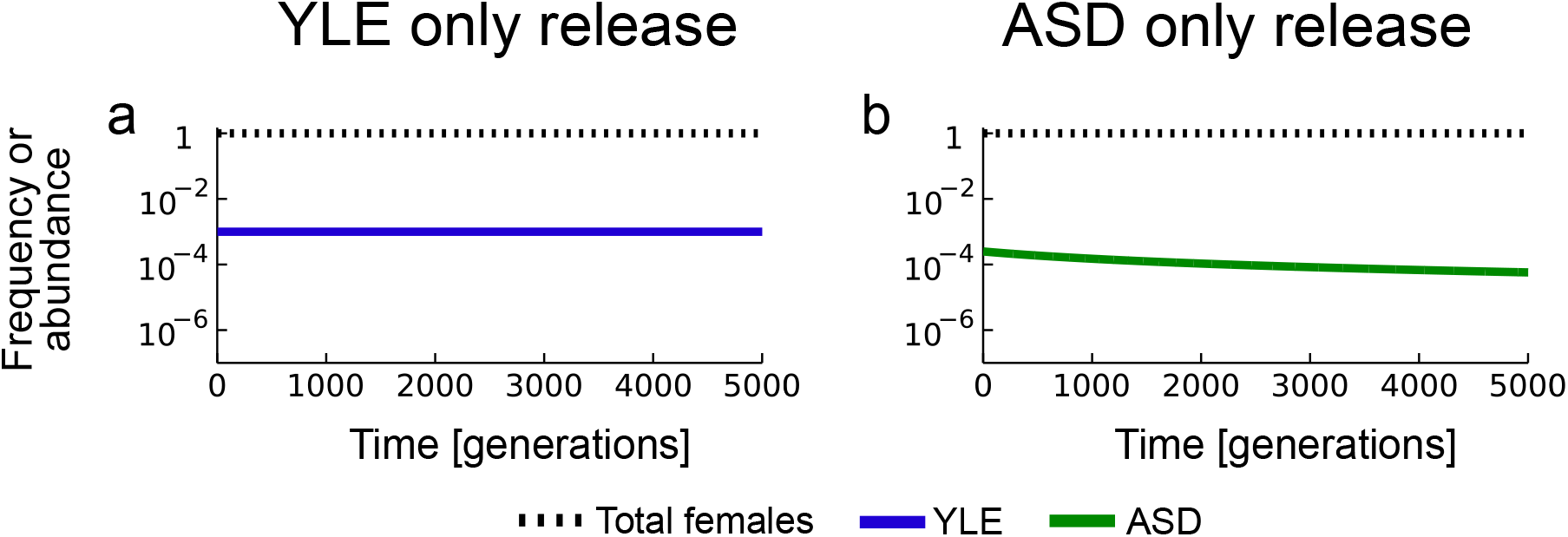
Timecourse of gene and population dynamics when only one construct is released, either the YLE (a) or the ASD (b) for the idealised case of no mutation, resistance, or unintended fitness costs.

**Supplementary Figure SF-2:**
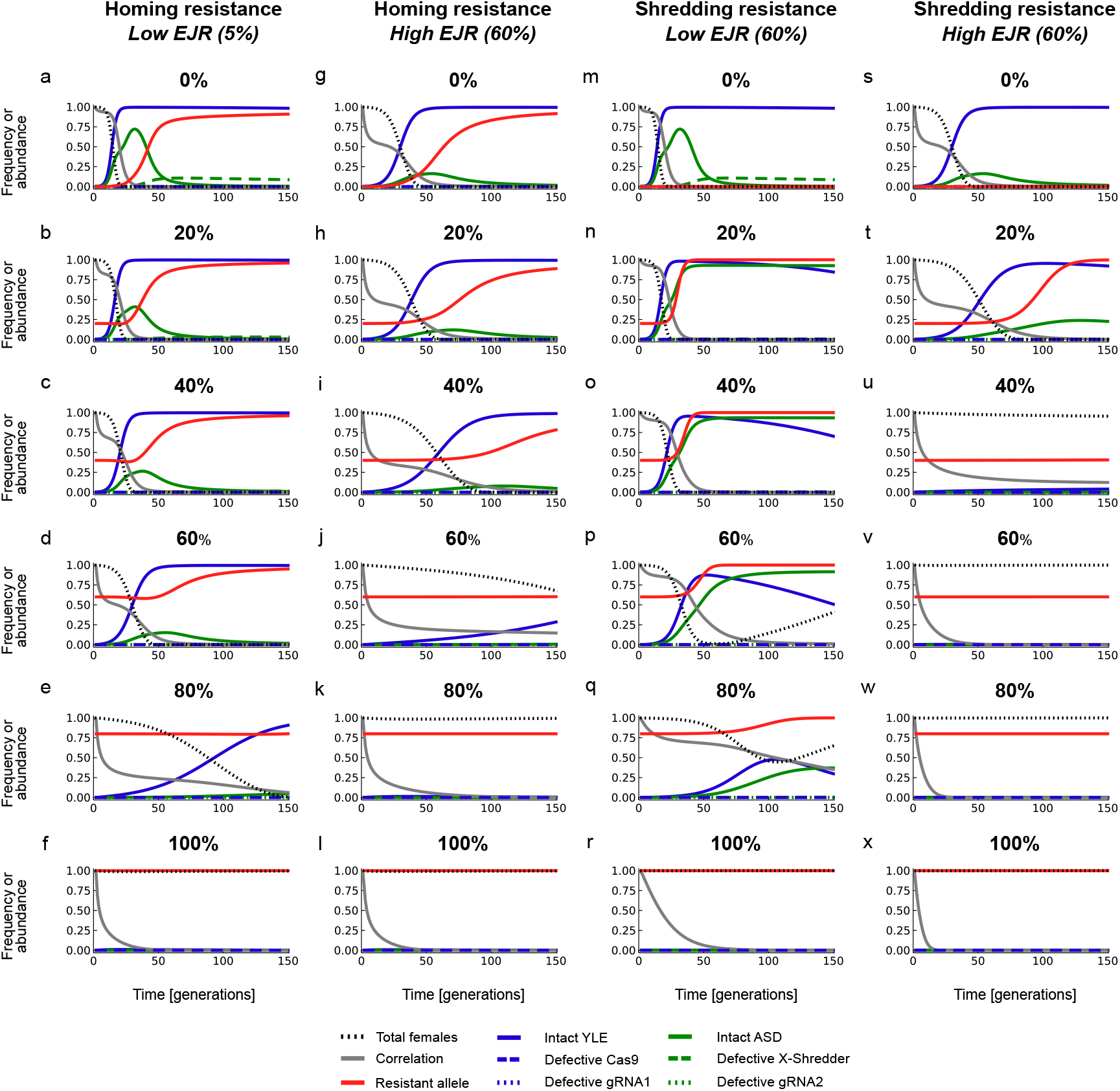
Timecourses for gene and population dynamics different initial frequencies (0, 20, 40, 60, 80 or 100%) of either homing or shredding resistance in low and high EJR species.

## Supplementary information: the model and PAM site analysis

### Model organism and ecology

The model considers an infinite population with discrete non-overlapping generations, separate sexes and male heterogamety. Mating is random, and all females mate (i.e., males are not limiting). There are two life stages, juveniles and adults, and density-dependent and -independent mortality occur at the juvenile stage, with the chance of surviving to adulthood being equal to Θ ∗ α / (α + N_h_[t]), where Θ represents non-specific density-dependent mortality, α the strength of density-dependent mortality, and N_h_[t] the total number of juveniles at that time point. A given intrinsic rate of increase of the population (*R*_*m*_) is used to calculate the number of eggs produced per female.

Transgenic releases are assumed to be performed as releases of adult males that have equal mating success as wild males.

### Molecular implementation

While the proposed strategy can be implemented in a number of ways, the specific genetic instantiation we model consists of four different genetic elements (Fig. 1 in main text). We assume the YLE to be constructed using a CRISPR/Cas type RNA-guided endonuclease system made up of a nuclease component, i.e. Cas9, and a guide RNA (gRNA1) component that allows editing of the target gene. Further, we assume the autosomal element consists of an X-shredder nuclease and a second guide RNA (gRNA2) that allows the autosomal construct to home in the presence of the YLE. All components can independently become defective due to mutation.

In total, the model considers four different loci: a locus on the Y chromosome where the YLE would be inserted, an autosomal locus where the other construct would be inserted, a locus on the X-chromosome that is the target of editing, and another X-linked site that is the target of X-shredding. The last one may consist of an array of tandem repeats, but for simplicity we model it as a single locus. Given that each of the components of the insertions can acquire loss-of-function mutations, and each of the target sites can acquire resistant mutations, there are 5 different Y chromosome variants, 6 autosomal variants and 7 X chromosome variants (Supplementary Figure SF-3). All biologically permitted combinations of chromosomes lead to 1071 different genotypes being considered in the model.

**Supplementary Figure SF-3:**
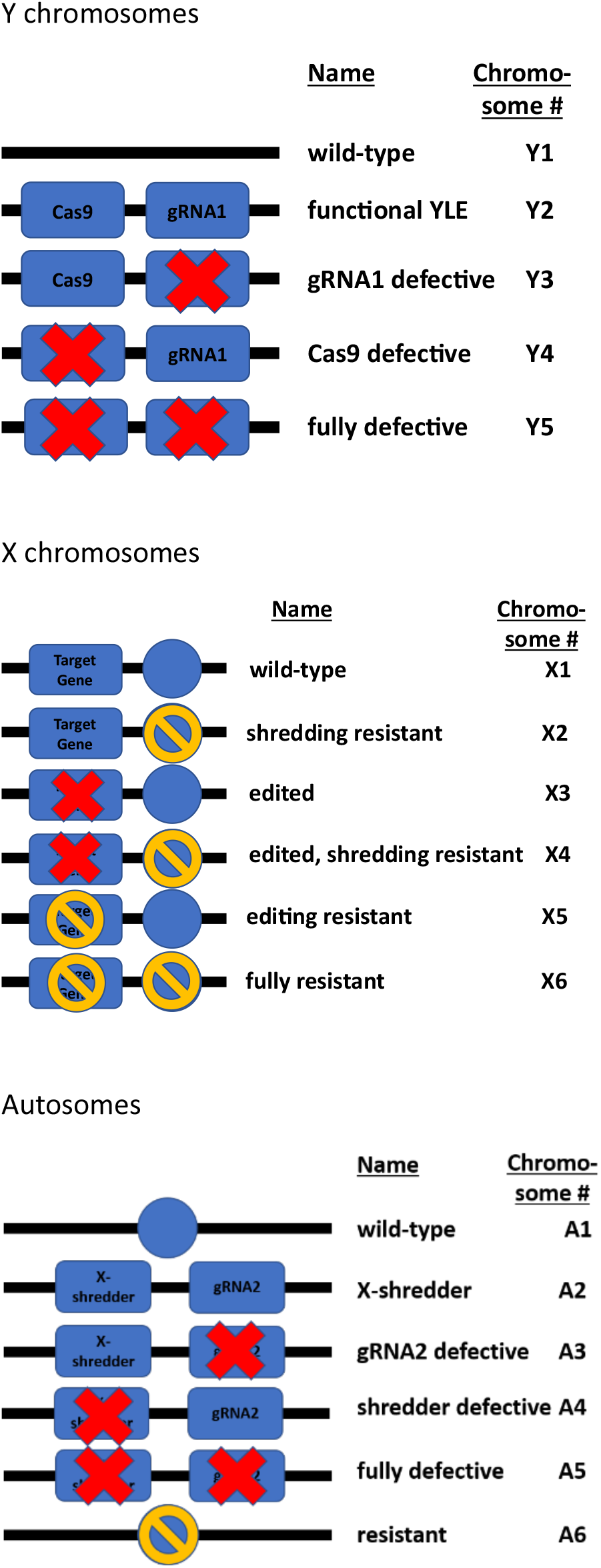
Overview of the different chromosome variants in the model

### Editing

Editing is assumed to occur in the germline of males carrying a functional YLE and an editable wild-type target gene. The frequency with which editing effectively occurs is described by the parameter *e*_*e*_ (efficiency of editing). While the intention of the editing is to produce a non-functional or dominant negative allele, occasionally, with probability *e*_*r1*_, a functional allele that is equally fit as the wildtype and is resistant to further editing is produced (Supplementary Figure SF-4). The resulting editing-resistant variant is assumed to have equal (i.e., unaffected) fitness as the wild-type.

**Supplementary Figure SF-4:**
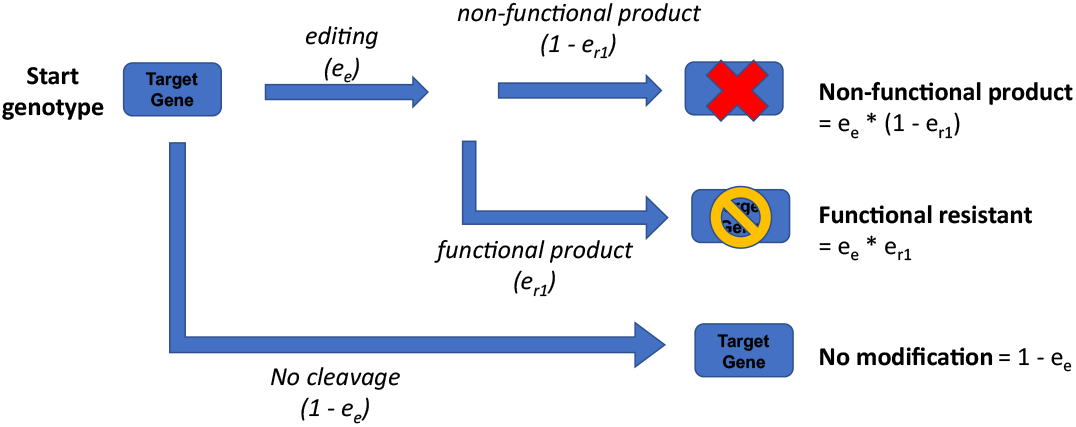
Schematic representation of the editing process.

### Homing

Homing is assumed to occur when a functional Cas9, a functional gRNA2 and a wild-type autosome occur in the same individual. The homing process begins with cleavage of the wild-type allele, which occurs with probability *e*_*h*_. This is then repaired by end joining, creating a homing-resistant allele (probability *e*_*r3*_), or by homology-directed repair (probability 1-*e*_*r3*_). When there is homology-directed repair, the gRNA2 and the X-shredder (if it exists) may each acquire loss-of-function mutations during the homing process with probability *m*_*1*_ (Supplementary Figure SF-5).

**Supplementary Figure SF-5:**
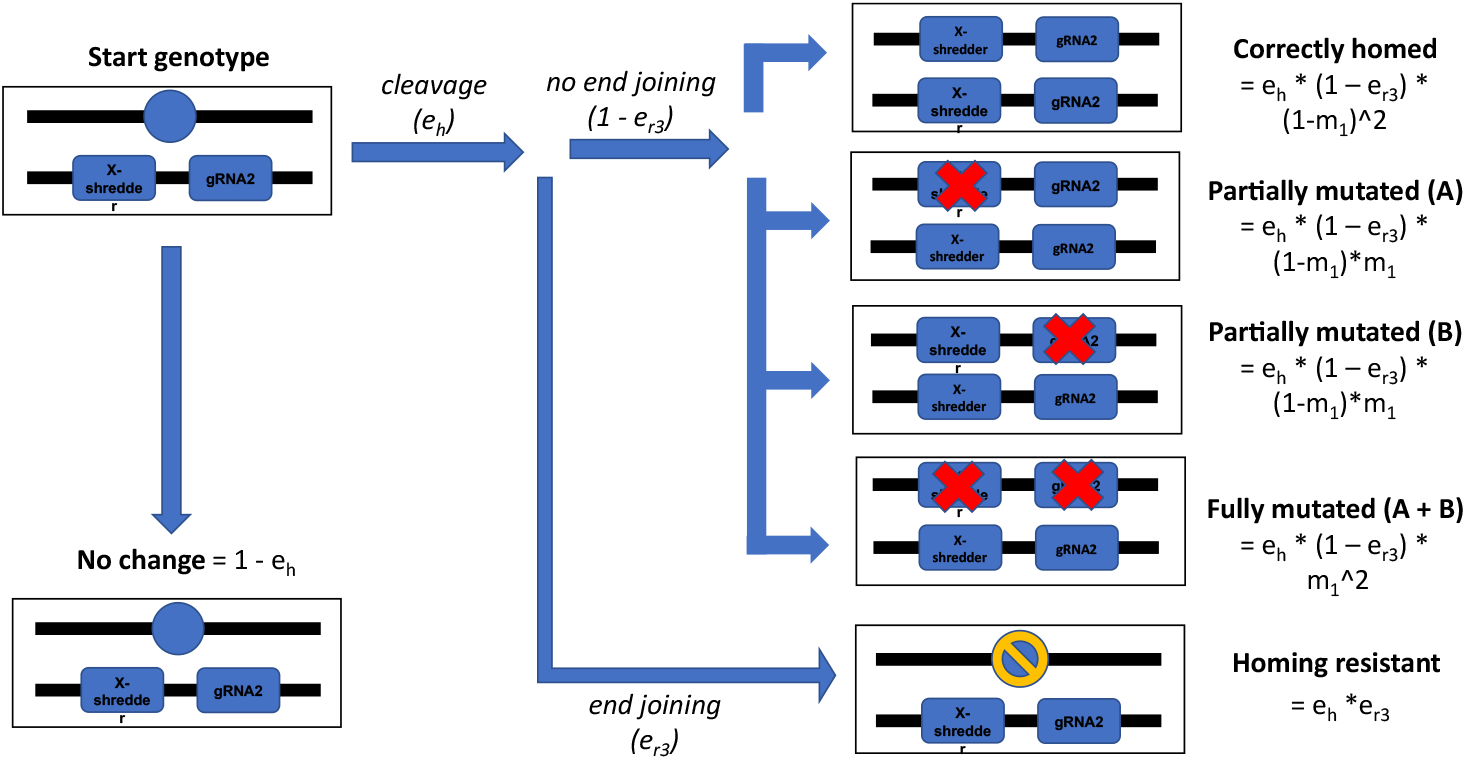
Schematic representation of the homing process.

### Shredding

Shredding is assumed to occur during sperm production and distorts the frequency with which sex chromosomes are passed on. Parameter *e*_*s*_ *(efficiency of shredding)* describes how frequently X chromosomes are shredded in X-shredder homozygotes, with *h*_*e2*_ being its dominance coefficient for X-shredder heterozygotes. We assume throughout that the shredding rate is not dosage dependent (*h*_*e2*_=1). Shredded X-chromosomes are repaired and resistant to further shredding with probability *e*_*r2*_.

### Recombination

Recombination may occur, with probability *r*, between the X-linked targes of editing and shredding. It only has an effect in females that are double heterozygotes.

### Mutation

All transgenic elements become defective through loss-of-function mutations during normal DNA replication with probability *m*_*2*_.

### Selection

The fitness of wildtype genotypes is standardised to 1. For simplicity, all types of selection via imposed fitness costs are assumed to occur at the same time, via differential mortality at the adult stage (i.e., after density dependent larval mortality). Many different factors can reduce fitness.

Parameter *s*_*f*_ describes the fitness cost for females homozygous for the edited target gene, *h*_*f*_ the corresponding dominance factor of that fitness cost, and *s*_*m*_ is the fitness cost for males carrying an edited target gene. Fitness costs of the different transgenic elements are broken down into different categories to allow for partially defective variants. Parameter *s*_*a*_ represents the expression cost of the Cas9 protein, *s*_*b*_ the expression cost of a gRNA, and *s*_*c*_ the expression cost of the X-shredder protein. Further, we suppose there is an additional activity associated cost when Cas9 is in the presence of a gRNA (*s*_*d*_) and an activity cost of the X-shredder (*s*_*e*_), which has dominance coefficient (*h*_*e*_) in heterozygous individuals. We assume throughout that the activity cost of X-shredding is not dosage dependent (*h*_*e*_=1).

### The order of events

The order of events is as shown in Supplementary Figure SF-6. All plots in the paper show the state of the population as censused at the 9th stage, showing somatic genotypes.

**Supplementary Figure SF-6:**
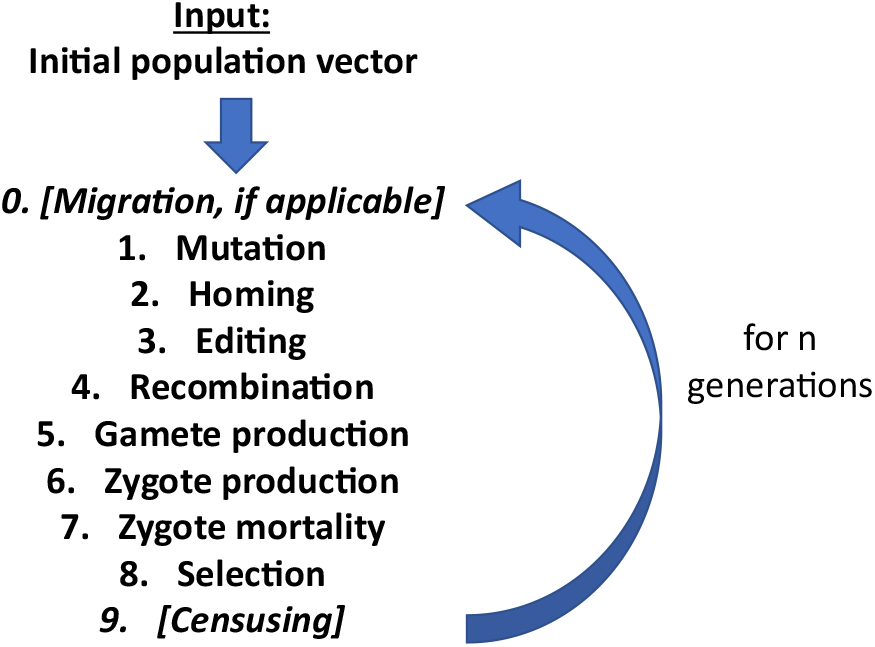
The order in which different life cycle processes were modelled

### Baseline parameter values

A list of parameters in the model and their baseline values for the idealised case and the sensitivity analyses in the main paper are shown in Supplemental Table S-1.

**Supplemental Table S-1:** List of parameters and their baseline values.

| Parameter name | Description | Ideal baseline value | Sensitivity scan baseline value |
| --- | --- | --- | --- |
| **Molecular efficiencies** |  |  |  |
| $e_e$ | efficiency of editing an X-linked target gene | 0.95 | 0.95 |
| $e_s$ | efficiency of X-shredding | 0.90 | 0.90 |
| $e_h$ | efficiency of cleaving the autosomal homing site | 0.95 | 0.95 |
| **Fitness effects** |  |  |  |
| $s_f$ | selection coefficient against homozygous females carrying the edited X-linked gene | 1 | 1 |
| $h_f$ | dominance coefficient for heterozygous females carrying only one edited allele | 1 | 1 |
| $s_a$ | fitness cost of Cas9 expression | 0 | 0 |
| $s_b$ | fitness cost gRNA1 / gRNA2 expression | 0 | 0 |
| $s_c$ | fitness cost of X-shredder expression | 0 | 0 |
| $s_d$ | fitness cost of Cas9-gRNA complex activity | 0 | 0.01 |
| $s_e$ | fitness cost of X-shredder activity | 0 | 0.01 |
| $s_m$ | selection coefficient against males carrying an edited target gene | 0 | 0 |
| **Loss-of-function mutation rates** |  |  |  |
| $m_1$ | mutation rate during homing | 0 | 1e-3 |
| $m_2$ | background mutation rate | 0 | 1e-6 |
| **Probabilities of resistance arising** |  |  |  |
| $e_{r1}$ | probability the edited allele is functional and resistant to further editing | 0 | 0 |
| $e_{r2}$ | probability a shredded X-chromosome is repaired and resistant to further shredding | 0 | 0 |
| $e_{r3}$ | probability the cleaved target of homing is repaired by end-joining | 0 | 0.05, 0.6 |
| **Miscellaneous** |  |  |  |
| $r$ | recombination rate of X-linked targets of editing and shredding | 0.5 | 0.5 |
| $R_m$ | intrinsic rate of population growth | 6 | 6 |

### PAM site analysis

Our analyses used population genomic sequence data from 1138 individuals from 15 populations, including *An. gambiae* ss, *An. coluzzii*, and populations of uncertain assignment, since gene flow is known to occur between them. The populations and sample sizes are as follows: *An. gambiae* from Cameroon (594), Uganda (224), Burkina Faso (184), Gabon (138), Guinea (80), Mayotte (48), Ghana (24), Bioko (18); *An coluzzii* from Angola (156), Burkina Faso (150), Cote d’Ivoire (142), Ghana (110); and populations of uncertain assignment from Guinea-Bissau (182), The Gambia (130) and Kenya (96) where n is the number of sequences and n/2 is the number of individuals sampled.

The frequencies of potential PAM sites (i.e., GG and CC dinucleotides) were calculated for each population, and sites with more than 5% missing data within any population were removed, leaving 13,462,450 polymorphic PAMs. For each population pair we counted the number of PAM sites which were present in one population at >36% frequency and in the other at <13%, representing the level of differentiation required at the homing target site in low EJR species to limit impact to the first population, then reversed the assignment of populations (as if the second was the target), and then averaged the two counts. The analysis was then repeated using the thresholds appropriate for high EJR species (>88% in one population and <33% in the other). F_ST_ values were obtained from *Anopheles gambiae* Genomes Consortium. 2020: Supp fig S5. The sample site map was generated using the cartopy python package.

## Footnotes

1 Numbers given for specificity – ‘low’ somewhat meaningless.

## Notes

### Competing Interest Statement

The authors have declared no competing interest.

